# Characterizing the serotonin biosynthesis pathway upon aphid infestation in *Setaria viridis* leaves

**DOI:** 10.1101/642041

**Authors:** Anuma Dangol, Beery Yaakov, Georg Jander, Susan R Strickler, Vered Tzin

## Abstract

*Setaria viridis* (green foxtail millet), a short life-cycle C4 plant in the Gramineae, serves as a resilient crop that provides good yield even in dry and marginal land. Although *S. viridis* has been studied extensively in the last decade, its defense responses, in particular the chemical defensive metabolites that protect it against insect herbivory, are unstudied. To characterize *S. viridis* defense responses, we conducted transcriptomic and metabolomic assays of plants infested with aphids and caterpillars. Pathway enrichment analysis indicated massive transcriptomic changes that involve genes from amino acid biosynthesis and degradation, secondary metabolites and phytohormone biosynthesis. The Trp-derived metabolite serotonin was notably induced by insect feeding. Through comparisons with known rice serotonin biosynthetic genes, we identified several predicted *S. viridis* Trp decarboxylases and cytochrome P450 genes that were up-regulated in response to insect feeding. The function of one Trp decarboxylase was validated by ectopic expression and detection of tryptamine accumulation in *Nicotiana tabacum*. To validate the defensive properties of serotonin, we used an artificial diet assay to show reduced *Rhopalosiphum padi* aphid survival with increasing serotonin concentrations. This demonstrated that serotonin is a defensive metabolite in *S. viridis* and is fundamental for understanding the adaptation of it to biotic stresses.

**HIGHLIGHT:** A combined transcriptomic and metabolomic profiling of *Setaria viridis* leaves response to aphid and caterpillar infestation identifies the genes related to the biosynthesis of serotonin and their function in defense.

## INTRODUCTION

In nature, plants are continuously challenged by diverse insect herbivores. In response to insect infestation, plants produce constitutive and inducible defenses to reduce damage and enhance their own fitness (Zhou *et al.*, 2015). Although many plant defenses are produced constitutively during a specific developmental stage, regardless of insect attack, others are inducible in response to insect damage. Examples of herbivore-induced defense strategies are the accumulation of chemical defenses such as benzoxazinoids and glucosinolates, physical barriers such as the increased density of thorns, spikes or glandular trichomes, as well as the synthesis of protease inhibitors and proteases (Bennett and Wallsgrove, 1994; Boughton *et al.*, 2005; van Loon *et al.*, 2006; Ahuja *et al.*, 2010; Markovich *et al.*, 2013). These inducible defenses usually are present at low levels and become more abundant in response to insect feeding. The general herbivore-induced defense strategies are mediated by signaling pathways, including those mediated by jasmonic acid and salicylic acid (Kerchev *et al.*, 2012), which allow plants to conserve metabolic resources and energy for growth and reproduction in the absence of insect herbivory.

The aphid family (order Hemiptera, family Aphididae), which comprises approximately 5,000 species distributed worldwide (Rebijith *et al.*, 2017), causes massive yield losses due to both direct and indirect crop damage (Guerrieri and Digilio, 2008). Aphids consume water and nutrients from plants, while transmitting toxins through their saliva (Rabbinge *et al.*, 1981; Bing *et al.*, 1991; Zhou *et al.*, 2015; Tzin *et al.*, 2015). Aphids also are responsible for transmission of 40% of all plant viruses, including the most harmful of plant viruses (Fereres *et al.*, 1989; Nault, 1997). Another major pest for graminaceous plants are leaf-chewing insects such as lepidopteran larvae. In response to the lepidopteran attack, plants massively modify their transcriptome and synthesize chemical defense compounds (Oikawa *et al.*, 2004; Niemeyer, 2009; Glauser *et al.*, 2011). Many plant deterrent compounds are derived from catabolism of the aromatic amino acids Trp (tryptophan), Tyr (tyrosine) and Phe (phenylalanine) (Zhou *et al.*, 2015; Tzin *et al.*, 2015, 2017; Wisecaver *et al.*, 2017). In the Gramineae family, indole, and its precursor Trp, serve as sources for several classes of defensive secondary metabolites including i) benzoxazinoids in maize (*Zea mays*), wheat (*Triticum aestivum*), rye (*Secale cereal)*, and wild barley species (Grün *et al.*, 2005; Ishihara *et al.*, 2017; Niculaes *et al.*, 2018; Zhou *et al.*, 2018), ii) gramine in cultivated barley (Grün *et al.*, 2005), and iii) serotonin (5-hydroxytryptamine) in rice (*Oryza sativa*), *Setaria italica* (foxtail millet), and *Echinochloa esculenta* (Japanese barnyard millet) (Ishihara *et al.*, 2008*b*, 2017; Lu *et al.*, 2018) (Fig. 1).

**Fig. 1.**
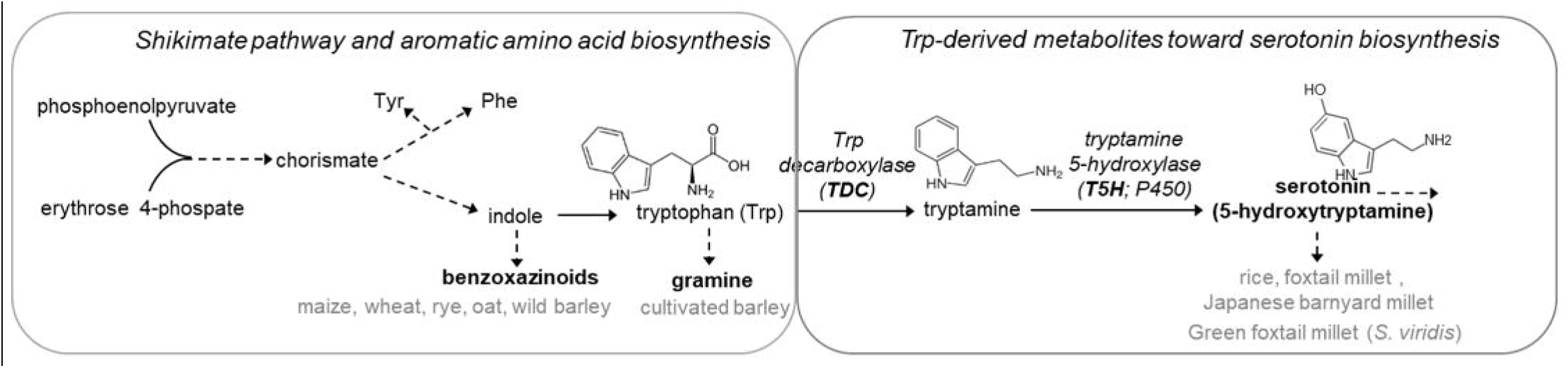
A schematic representation of the shikimate pathway, the aromatic amino acids, and synthesis of Trp-derived defensive metabolites. In italics font are the names of the enzymatic steps. In bold font are the major classes of defense metabolites. In gray are several plant species from the Gramineae family that produce this class of defense metabolites. Dashed arrows indicate multiple enzymatic steps.

Serotonin, a well-studied neurotransmitter in mammals, also has been found at detectable levels in more than 90 plant species (Kang *et al.*, 2008; Erland *et al.*, 2016; Alexandra *et al.*, 2017). In plants, it has been implicated in adaptations to environmental changes and in defense responses against pathogen infection and insect herbivory (Ishihara *et al.*, 2008*b*; Kang *et al.*, 2008; Park *et al.*, 2009). Serotonin biosynthesis requires the sequential action of two enzymes: first Trp decarboxylase (TDC), which produces tryptamine (Gill *et al.*, 2003; Ishihara *et al.*, 2011; Li *et al.*, 2016; Welford *et al.*, 2016), and second, tryptamine 5-hydroxylase (T5H), a cytochrome P450 that catalyzes the conversion of tryptamine to serotonin (Fujiwara *et al.*, 2010; Dharmawardhana *et al.*, 2013; Lu *et al.*, 2018) (Fig. 1). Both enzymes play an important role in determining serotonin levels and are induced by both chewing insect and pathogen attacks (Kang *et al.*, 2009; Hayashi *et al.*, 2015). Still, the function of serotonin in plants is inconclusive (Alexandra *et al.*, 2017). For example, ectopic expression of the *Camptotheca acuminata TDC1* gene allowed sufficient tryptamine to accumulate in poplar and tobacco leaf tissues to significantly suppress the growth of two moth species *Malacosoma disstria* and *Manduca sexta*, respectively (Gill *et al.*, 2003; Gill and Ellis, 2006). Similarly, ectopic expression of the *Aegilops variabilis TDC1* gene in tobacco plants increased resistance to the cereal cyst nematode, *Heterodera avenae* (Huang *et al.*, 2018). On the other hand, a recent study of the mutated *CYP71A1* (T5H) in rice, demonstrated that reduced serotonin levels caused higher susceptibility to rice blast *Magnaporthe grisea*, but also conferred resistance to rice brown spot disease (*Bipolaris oryzae*), and the rice brown planthopper (*Nilaparvata lugens*) (Lu *et al.*, 2018). The function of serotonin and tryptamine in defense against aphids has not yet been studied.

The Gramineae, a large plant family consisting of approximately 12,000 species and 771 genera (Soreng *et al.*, 2015), includes staple crops such as rice, wheat, maize, barley, sorghum and several millets (Brutnell *et al.*, 2010; Soreng *et al.*, 2015; Kokubo *et al.*, 2016*a*; Doust and Diao, 2017). *Setaria*, which serves as both a research model and a crop plant is a genus of panicoid C4 grasses (Brutnell, 2015; Saha and Blumwald, 2016), closely related to grasses such as switchgrass (*Panicum virgatum*) and pearl millet (*Panicum glaucum*), as well as to maize and sorghum (Li and Brutnell, 2011; Pant *et al.*, 2016). The domesticated *S. italica* is an ancient cereal grain crop from China, which expanded to India and Africa, excelling as a drought- and low-nutrient-tolerant grain (Goron and Raizada, 2015; Nadeem *et al.*, 2018), while its wild ancestor, *Setaria viridis* (green foxtail millet), is a widespread weed across the globe (Hu *et al.*, 2018). Both *S. italica* and, *S. viridis* genotypes have recently been sequenced, and their genomes are publicly available (Bennetzen *et al.*, 2012). *Setaria* species have thus recently emerged as model plants for studying C4 grass biology and other agronomic traits (Brutnell *et al.*, 2010). *Setaria viridis* has many advantageous properties as a monocot genetic model system: i) a short generation time of 6-8 months (illustrated in Fig. 2A); ii) a small sequenced diploid genome containing approximately 395 Mb and 38,000 protein-coding genes; iii) transient and stable transformation systems and established protocols; and iv) a small stature and simple growth requirements (Li and Brutnell, 2011; Liu *et al.*, 2016; Mei *et al.*, 2016; Van Eck *et al.*, 2017; Zhu *et al.*, 2018). Recent studies have exploited these attributes to achieve a comprehensive understanding of the dynamic gene expression of multiple inflorescence stages (Zhu *et al.*, 2018) and different tissues and stages of development, under abiotic stresses (Martins *et al.*, 2016; Saha *et al.*, 2016).

**Fig. 2.**
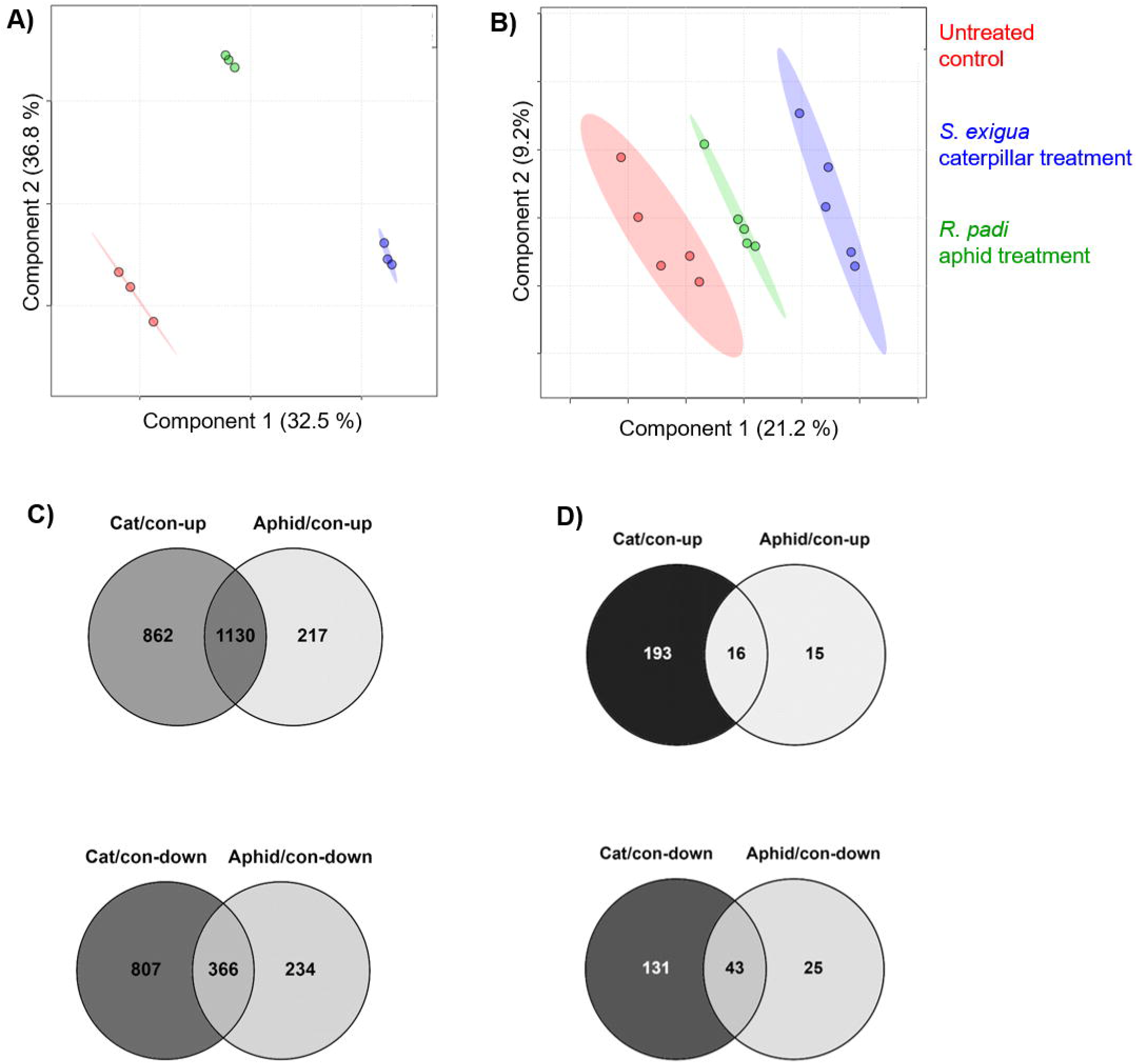
Overview of *Setaria viridis* transcriptome and metabolome responses to aphid and caterpillar feeding. A) PLS-DA plot of 15,000 genes identified by transcript profiling (RNA-seq) of *S. viridis* accession A10.1 infested with aphids and caterpillars. B) PLS-DA of 3,323 mass signals (negative and positive ion modes) identified by LC-TOF-MS. Ovals indicate 95% confidence intervals. C) and D) Venn diagrams illustrating the number of individual transcripts and mass spectrometry features that were significantly up- or down-regulated by each treatment, respective to untreated control. *P* value < 0.05, and fold change greater than 2 or less than 0.5.

In this research, we integrated transcriptomic and metabolomic datasets to identify changes that occur in response to herbivore feeding. We identified *S. viridis* genes that are involved in serotonin biosynthesis and suggest that this pathway plays a role in plant defense against herbivores. By the ectopic expression of a Trp-decarboxylase gene (*TDC1*), we demonstrated that the encoded enzyme converts Trp into tryptamine. By exposing aphids to an artificial diet supplemented with different concentrations of serotonin and tryptamine, we were able to demonstrate toxic functions of these molecules relative to Trp. Overall, our results indicate that serotonin biosynthesis is triggered by aphid feeding and that it is a deterrent metabolite in *S. viridis* leaves.

## MATERIALS AND METHODS

### Plant growth and insect bioassays

The *Setaria viridis* A10.1 accession (Brutnell *et al.*, 2010; Huang *et al.*, 2014) was used for all experiments. Plants grew for 15 days in a growth room under a controlled photoperiod regime with a 16-h light/8-h dark cycle, at a constant temperature of 25 °C and 60% relative humidity. The *Rhopalosiphum maidis* (corn leaf aphid) aphid was reared on B73 maize seedlings as described previously (Meihls *et al.*, 2013). The *Rhopalosiphum padi* (bird cherry-oat aphid) was reared on barley seedlings (Noga cultivar, Negev Seeds, Israel). For the *Spodoptera exigua* (beet armyworm) eggs were purchased from Benzon Research (Carlisle, PA, USA), and kept for 48 h in a 29 °C incubator. After hatching, first instar caterpillars were transferred to an artificial diet. For transcriptomic analysis, ten adult *R. maidis* aphids or three 2^nd^-3^rd^ instar *S. exigua* were placed on 15-day old *S. viridis* plants for 96 h (five plants for each replicate), in addition to untreated control plants, and covered with micro-perforated polypropylene bags (whole cage). To identify defense metabolites and confirm the gene expression by using qRT-PCR, 15-day old *S. viridis* plants were subjected to infestation with ten adult *R. padi* in a whole cage setup for 6, 24, 48 and 96 h, in addition to the untreated control (0 h).

### Transcriptome sequencing and RNA-seq data analysis

Tissues from 4-6 plants were combined into one replicate, and four replicates were collected for each treatment (aphid, caterpillar, and untreated control). Total RNA was extracted using an SV Total RNA Isolation Kit with on-column DNase treatment (Promega) (Tzin *et al.*, 2015). For transcriptomic analysis, strand-specific RNA-seq libraries were prepared (Zhong *et al.*, 2011; Chen *et al.*, 2012). The purified RNA-seq libraries were quantified, and 20 ng of each were used for next-generation sequencing using an Illumina HiSeq2000 instrument at Weill Cornell Medical School with a 101 bp single-end read length. For RNA-seq read quality, values were checked using FASTQC. Then, by using fastq-mcf (https://github.com/ExpressionAnalysis/ea-utils/blob/wiki/FastqMcf.md), the adapters and low-quality sequences were trimmed and removed with a minimum length of 50 bp and a minimum quality value of 30 (Tzin *et al.*, 2017). The RNA-seq data were mapped to the *Setaria viridis* v2.1(Green foxtail) genome reference (https://phytozome.jgi.doe.gov/), using Hisat2 (2.0.5) (default parameters) (Kim *et al.*, 2015), and the resulting BAM files were filtered for hits with single-locus, concordant alignments of paired reads of sufficiently high mapping quality (MQ > 20). Expression levels of all the genes (including chromosome ‘Un’) were calculated using the DESeq2 package (Love *et al.*, 2014). We used CPM (count per million fragments mapped) values for quantification of gene expression (Supplementary Table S1).

### Confirming gene expression using quantitative PCR

Total RNA was extracted using an RNeasy Plant Mini Kit (Qiagen, http://www.qiagen.com/). The RNA was treated with DNase I and then purified further using an RNeasy binding column. Single-strand cDNA was synthesized from 1 µg of total RNA using a Verso cDNA Synthesis Kit (Thermo Fisher Scientific). Quantitative PCR reactions were performed using SYBR Green Master Mix (Bio-Rad), according to the manufacturer’s protocol. Primers for the TDC1 gene (Sevir.6G066200) were designed using Primer 3 plus software with the following parameters: product size range of 100-120 bp, primer size of 20 to 25 bp, primer Tm of 57 to 63 □C, primer GC% of 45 to 65 % and product Tm of 60 □C. To quantitatively determine the steady-state levels of transcripts, the 2^−ΔΔCt;^ method (Livak and Schmittgen, 2001) was used. The expression of genes was normalized against *Setaria* Eukaryotic Initiation factor 4A (e*IF4A;* Sevir.4G254900) as an internal reference (housekeeping gene) (Saha *et al.*, 2016). Each real-time PCR sample was run in triplicate. For primer list used for the qRT-PCR see supplementary Table S2.

### Cloning of *TDC1* gene and Agrobacterium□mediated transformation

The full coding sequence of *TDC1* (Sevir.6G066200) was retrieved from Phytozome (https://phytozome.jgi.doe.gov) and cloned using GoldenBraid cloning vectors, including pUPD1 and p3α2, that were developed by Diego Orzaez (Sarrion-Perdigones *et al.*, 2014). An internal *Bsm*BI site was substituted by a synonymous base, a 6xHA tag that was fused to the 3’ and a *Bsm*BI site was added at both ends according to user guidelines (https://gbcloning.upv.es/). The fused gene sequence was synthesized and cloned into pUC57 (acquired from Hylabs, Israel) The 3’ overhang of the introduced *Bsm*BI site on the *TDC1* plasmid was mutated by site-directed mutagenesis PCR to make it compatible for cloning into the pUPD1 vector. The domestication vector, pUPD1, and the *TDC1* PCR product were separately digested with *Bsm*BI (Thermo Fisher Scientific) at 37 °C overnight following the manufacturer’s protocol. Digested products were separated on a 1% agarose gel, purified and ligated using T4 DNA ligase (Thermo Fisher Scientific). The ligated product was transformed into *E. coli* DH5α strain using the heat-shock method at 42 °C for 1 min, and selected on Ampicillin, isopropylthio-β-galactoside (IPTG) and 5-bromo-4-chloro-3-indolyl-β-D-galactopyranoside (X-gal). The positive colonies (white) were confirmed by PCR followed by sequencing. PUPD1 harboring *TDC1*, the *Arabidopsis thaliana Ubiquitin3* promoter, and THI4 terminator plasmids were assembled into destination vector p3α2 using *Bsa*I restriction-ligation reaction. The restriction-ligation reaction was set up by mixing 75 ng of each plasmid, 1 ul *Bsa*I, 1 ul T4 ligase, 2.5 ul 10x ligase buffer and 2.5 ug BSA. The reaction was incubated for 10 min at 37 °C and 100 cycles of (3 min at 37 °C and 4 min at 16 °C), followed by 10 min at 50°C and 10 min at 80 °C. The ligated product was transformed into *E. coli* DH5α competent cells as described above. Transformants were selected with Kanamycin, IPTG, and X-gal, and the clones with correct assembly were confirmed by restriction with *Hind*III. The primer sequences that were used for cloning are described in Supplementary Table S3.

### TDC1 protein expression in *Nicotiana benthamiana* plants and immunoblot analysis

For the transient expression of *TDC1*, a plasmid was transformed into *Agrobacterium tumefaciens* strain GV3101 by heat-shock. Agroinfiltration was performed as described previously (Wieland *et al.*, 2006). In brief, overnight grown bacterial cultures were centrifuged and the pellet was re-suspended in agroinfiltration medium including 10 mM of MES pH 5.6, 10 mM of MgCl_2_, and 200 μM of acetosyringone, to an optical density of 0.4 at 600 nm. The leaf infiltration was performed by mixing equal volumes of the corresponding bacterial suspensions. Inoculations were carried out by syringe-agroinfiltration in leaves of 4-5 week old *Nicotiana benthamiana* plants (growing conditions: 24 °C day/ 20°C night in a 16 h light/8 h dark cycle). For TDC1 protein expression, leaf tissue was collected 1-5 days post-infiltration and examined. To detect the fusion protein, total protein was extracted from samples of 100 mg (fresh weight) liquid nitrogen ground leaf tissue containing 20 mg Polyvinylpolypyrrolidone (PVPP) and 200 μl potassium phosphate buffer (25 mM), protease inhibitors (Complete, ROCHE) and 1 mM PMSF. Soluble protein extracts were separated from cell precipitates by centrifugation at 15,000 g for 10 min and boiled in protein sample buffer (Manela *et al.*, 2015). Ponceau staining and de-staining were performed using Ponceau dye (Sigma-Aldrich) according to the manufacturer’s instructions. The immunoblot was performed as previously described (Stepansky and Galili, 2003), using monoclonal anti-HA antibodies (Sigma-Aldrich) and Clarity Western ECL Substrate (Bio-rad) following the manufacturer’s instructions and visualized in a chemiluminescence imager (Chemi-DoC, Bio-rad). The protein activity was validated by using GC-MS as described below.

### Targeted metabolic analysis using gas chromatography-mass spectrometry (GC-MS)

Metabolites were extracted using 400 mg of ground frozen plant tissue mixed with solvents containing a ratio of methanol/water/chloroform of 55:23:22, v/v following the protocol as previously described (Rosental *et al.*, 2016). After phase separation, 300 μl of the top hydrophilic layer, was collected and dried in a vacuum. Samples were derivatized by adding 40 μl of 20 mg/ml metoxyaminehy-drochloride (Sigma-Aldrich) dissolved in pyridine following incubation for 2 h in an orbital shaker at 1000 rpm at 37 °C. Next, N-methyl-N-(trimethylsilyl) tri-fluoroacetamide (MSTFA) including standard mix (Alkanes) in a volume of 77 μl was added to each sample and incubated for 30 min in an orbital shaker at 37 °C. Finally, 1 μl of the sample was injected into the Agilent 5977B GC-MS instrument. For data acquisition, we used the Mass Hunter software, and the NIST mass spectral library normalized to 13-sorbitol (Sigma-Aldrich) as an internal standard. For calibration curves, we used a mix of authentic standards: shikimate, Trp, tryptamine, serotonin (Sigma-Aldrich). For a list of primary metabolites detected by the GC-MS see Supplementary Table S4.

### Untargeted metabolic analysis using liquid chromatography-mass spectrometry (LC-MS/MS)

Whole plant aerial tissue was collected after 96 h of *R. maidis* or *S. exigua* infestation as well as untreated plants. Frozen ground tissue was weighed and extraction solvent (methanol/water/formic acid, 70:29.9:0.1, v/v/v) was added to each sample (1:3 ratio) follow by brief vortexing, shaking for 40 min at 4 °C, and centrifugation for 5 min at 14,000 g. The samples were filtered with a 0.45 μm filter plate (EMD Millipore Corp., Billerica, MA, USA) by centrifuging at 2,000 g for 3 min; then the supernatant was diluted 1:9, and transferred to glass vial (Mijares *et al.*, 2013; Tzin *et al.*, 2017). For mass feature detection, liquid chromatography-tandem mass spectrometry (LC-MS/MS) analysis was performed on a Dionex UltiMate 3000 Rapid Separation LC System attached to 3000 Ultimate diode array detector attached to Thermo Q-Exactive mass spectrometer (Thermo Fisher Scientific). The samples were separated on a Titan C18 7.5 cm×2.1 mm×1.9 μm Supelco Analytical Column (Sigma-Aldrich), using the same parameters as previously described (Handrick *et al.*, 2016). Raw mass spectrometry data files were processed with XCMS (Smith *et al.*, 2006) and CAMERA (Kuhl *et al.*, 2012), software packages for R. Both negative and positive ionization data sets were transferred to Microsoft Excel and presented in Supplementary Table S5.

### Assessment of aphid performance on an artificial diet

A previously described artificial diet assay involving indole alkaloids from Gramineae and *R. maidis* (Corcuera, 1984) was modified to measure the effects of Trp, tryptamine, and serotonin on *R. padi* survival. Aphid artificial diet, composed of twenty different essential amino acids and sucrose was adapted from Prosser and Douglas (1992). The amino acids were dissolved in 50 ml of double distilled water and filtered by using a 45 µm filter. Twenty adult *R. padi* were added to 35 ml plastic cups, a sheet of Parafilm (Pechiney Packaging Company) was stretched across the 4.5 cm diameter opening. The artificial diet was diluted 1:1 and supplemented with 0.28, 0.56 or 1.1 mM of Trp, tryptamine or serotonin. Then, 100 μl of the artificial diet with supplements was pipetted onto a Parafilm. To keep the solution in place, another sheet of Parafilm was stretched across the top (Prosser and Douglas, 1992; Meihls *et al.*, 2013). The number of adult aphids was counted after 1-2 d on the control or supplemented diet.

### Statistical analyses

Data for partial least-squares discriminant analysis (PLS-DA) plots were normalized as follows: an average of each parameter was calculated across all samples (untreated, caterpillar and aphid infestation), and each individual parameter was divided by its own average and subjected to a Log_2_ value (Shavit *et al.*, 2018). The plots were drawn using MetaboAnalyst 3.0 software (Xia *et al.*, 2009). The Venn diagrams were designed using the Venny 2.1.0 drawing tool (http://bioinfogp.cnb.csic.es/). The pathway enrichment analysis was performed using MetGenMAP (Joung *et al.*, 2009). For the metabolic dataset, statistical comparisons for Student’s *t*-tests, and an analysis of variance (ANOVA) was conducted using JMP13 (www.jmp.com, SAS Institute, Cary, NC, USA).

## RESULTS

### Transcriptomic and metabolomic dataset analyses of *Setaria viridis* responses to caterpillar and aphid feeding

To investigate the global transcriptomic changes in response to caterpillar and aphid feeding, we performed whole-plant, non-choice insect bioassays using four 2^nd^-3^rd^ *Spodoptera exigua* instars or twenty *Rhopalosiphum maidis* aphids for 96 h of infestation. Mapping of RNA-seq data to predicted gene models (v2.1) showed approximately 15,000 transcripts (Supplemental Table S1). The transcript levels were normalized, and partial least-squares discriminant analysis (PLS-DA) was conducted (Fig. 2A). Biological replicates from each treatment, as well as the control, clustered together, highlighting the reproducibility of the experiment. The PLS-DA plot showed that samples from the aphid and caterpillar treatments clustered separately from the control samples, indicating that a substantial change in the transcriptome occurred due to aphid and caterpillar feeding (components 1 and 2 explained 69.3 % of the variance). The differences between the total up- and down-regulated transcripts were calculated for each treatment and are presented in Venn diagrams showing genes with significant expression differences of *P* value ≤ 0.05 (false discovery rate [FDR] adjusted) and at least 2-fold expression difference (Fig. 2C; data in log values Supplemental Table S1). A total of 2,209 genes were up-regulated, 217 by aphids, 862 by caterpillars and 1,130 by both insects. And a total of 1,407 genes were down-regulated, 234 by aphids, 807 by caterpillars and 366 by both insects. This suggested a common pattern of gene expression as well as a unique one for each individual insect. Only a small group of 11 genes were up-regulated by caterpillars and down-regulated by aphids.

To understand the metabolic changes which occur following herbivore infestation, the same leaf samples were subjected to untargeted liquid chromatography-tandem mass spectrometry (LC-MS/MS) in negative and positive ion modes (Supplemental Table S5). The PLS-DA clustering pattern of the untargeted metabolomics analysis (both negative and positive ion modes) showed that samples of each treatment were clustered separately from the control. However, the aphid-infested samples were located between the control and caterpillar-infested clusters, which indicates an intermediate effect on the aphid-infested plants (Fig. 2B). Mass features with significant differences (*P* value ≤ 0.05) and at least 2-fold changes relative to the controls were selected, resulting in a total of 99 and 383 mass features significantly altered by aphids and caterpillars, respectively. A total of 224 mass features were increased, 15 by aphids, 193 by caterpillars and 16 by both insects. Additionally, a total of 199 mass features were reduced, 25 by aphids, 131 by caterpillars and 43 by both insects (Fig. 2D). This too suggests a common pattern of gene expression and a unique one for each individual insect. The caterpillar feeding modified a larger number of transcripts and mass features than the aphids which suggests a stronger effect of caterpillar infestation. The similarity of the transcriptomic and metabolomic dataset clustering, suggests both overlap in plant responses to herbivore attack, as well as a unique pattern for each herbivore.

### Pathway enrichment of the transcriptome data

To elucidate the metabolic processes that are involved in each gene group shown in the Venn diagrams (Fig. 2C), over-representation pathway enrichment analysis was performed using MetGenMAP (Joung *et al.*, 2009) to compare to rice orthologues (LOC gene ID; table Supplemental Table S6). In Table 1 the significantly enriched pathways of up- and down-regulated genes are presented (Table 1A and 1B respectively). The pathways that were significantly enriched by both caterpillar and aphid feeding were mainly associated with amine and polyamine degradation, amino acid metabolism, and biosynthesis of carbohydrates, cell structures, fatty acids, lipids, hormones and secondary metabolites. The genes related to amino acid biosynthesis and their degradation into secondary metabolites of phenylpropanoids and salicylic acid and cell structures biosynthesis (suberin) were associated with caterpillar feeding. Additionally, genes related to secondary metabolites degradation (betanidin) and cofactor, prosthetic group, electron carrier biosynthesis (chlorophyllide a biosynthesis) were uniquely enhanced by aphid feeding. As presented in Table 1B, only a few pathways were significantly enriched among the down-regulated genes by both insects, including the degradation of amino acids, Tyr, Arg and beta-alanine, and citrulline biosynthesis as well as coenzyme A and D-lactate fermentation. Down-regulated genes from the amino acids Val, Ile, Leu and Thr were enriched by caterpillar feeding as well as cofactors, prosthetic groups, electron carrier biosynthesis, Acyl-CoA thioesterase, and triacylglycerol degradation and similar pathways were enriched by aphid feeding. No significantly altered pathways were found in the group significantly up-regulated by caterpillars and down-regulated by aphids (Table 1C). Together, these observations indicate that there is a shift in primary metabolism processes including amino acids, carbohydrates, and fatty acids and lipids and the production of secondary metabolites, cell structures and phytohormones in response to aphid and caterpillar feeding on *S. viridis*.

**Table 1.**
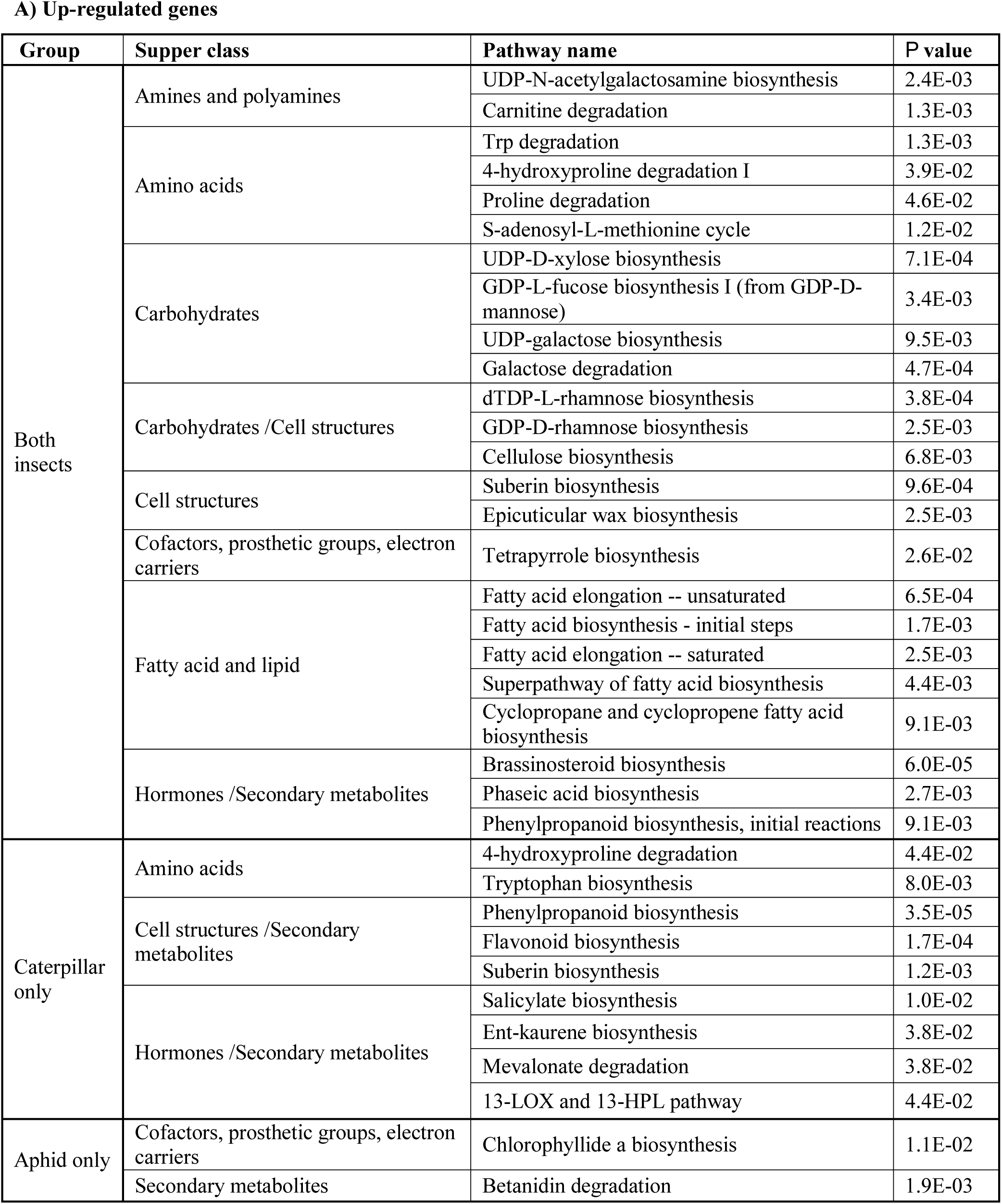
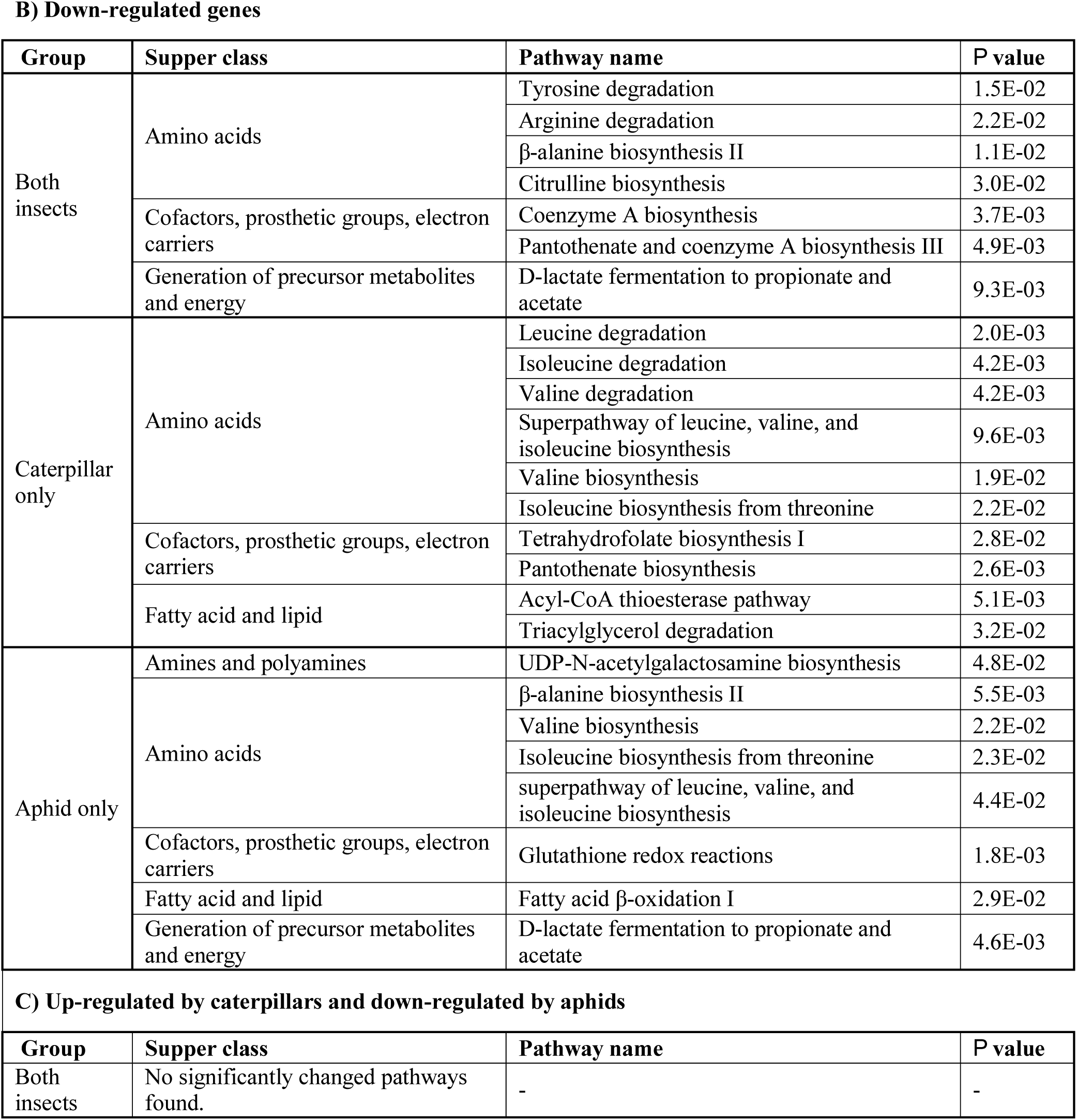
Pathway enrichment analysis of significantly altered gene expression in response to caterpillar and aphid feeding. The genes were classified into seven groups: A) Up-regulated genes by both insects, caterpillar only, and aphid only feeding; B) Down-regulated genes by both insects, caterpillar only, or aphid only feeding; C) Genes that were up-regulated by caterpillars and down-regulated by aphids. The enrichment analysis and pathway classification was performed using MetGenMAP (*P* value <0.05) (Joung *et al.*, 2009), using the rice orthologues. Pathway name indicates the specific pathway, and supper class indicates the general metabolic classification.

### Plant hormone-related genes induced by aphid and caterpillar feeding

The overrepresented pathway enrichment suggested that *S. viridis* defense signaling pathways are differentially regulated by aphid and caterpillar feeding (Table 1). To identify the transcriptional signatures of hormonal responses in herbivore-infested *S. viridis* plants, we used the Hormonometer program, which compares the variation in gene expression with indexed data sets of hormone treatments (Oa *et al.*, 2009). We evaluated the similarity in gene expression profiles elicited by aphids and caterpillars to the application of the following plant hormones: jasmonic acid (methyl jasmonate), 1-aminocyclopropane-1-carboxylic acid (a precursor of ethylene), abscisic acid, auxin, cytokinin (zeatin), brassinosteroid, gibberellin (GA3) and salicylic acid. Because the hormone treatments were conducted with *Arabidopsis thaliana*, we selected the *S. viridis* orthologous genes from *Arabidopsis* containing probeset identifiers, which are required by Hormonometer (a total of 9,017 orthologous genes were used; Supplemental Table S7). As presented in Fig. 3, the most common insect-induced *S. viridis* gene expression changes were associated with jasmonic acid, abscisic acid and auxin-dependent signaling. Salicylic acid-responsive genes were induced by both insects, with higher induction by aphids. Also, there was an overall positive correlation between herbivore-induced genes and those that were induced within 60 and 180 min of cytokinin and brassinosteroid-dependent signaling and 39 h of gibberellin treatment. Ethylene-responsive genes showed a general pattern of negative correlation with herbivore feeding from *S. viridis* leaves, except for the first treatment of 30 min that was positive in aphid feeding. Overall, the hormone patterns induced by aphids and caterpillars were similar, with some differences in the time of hormone treatments.

**Fig. 3.**
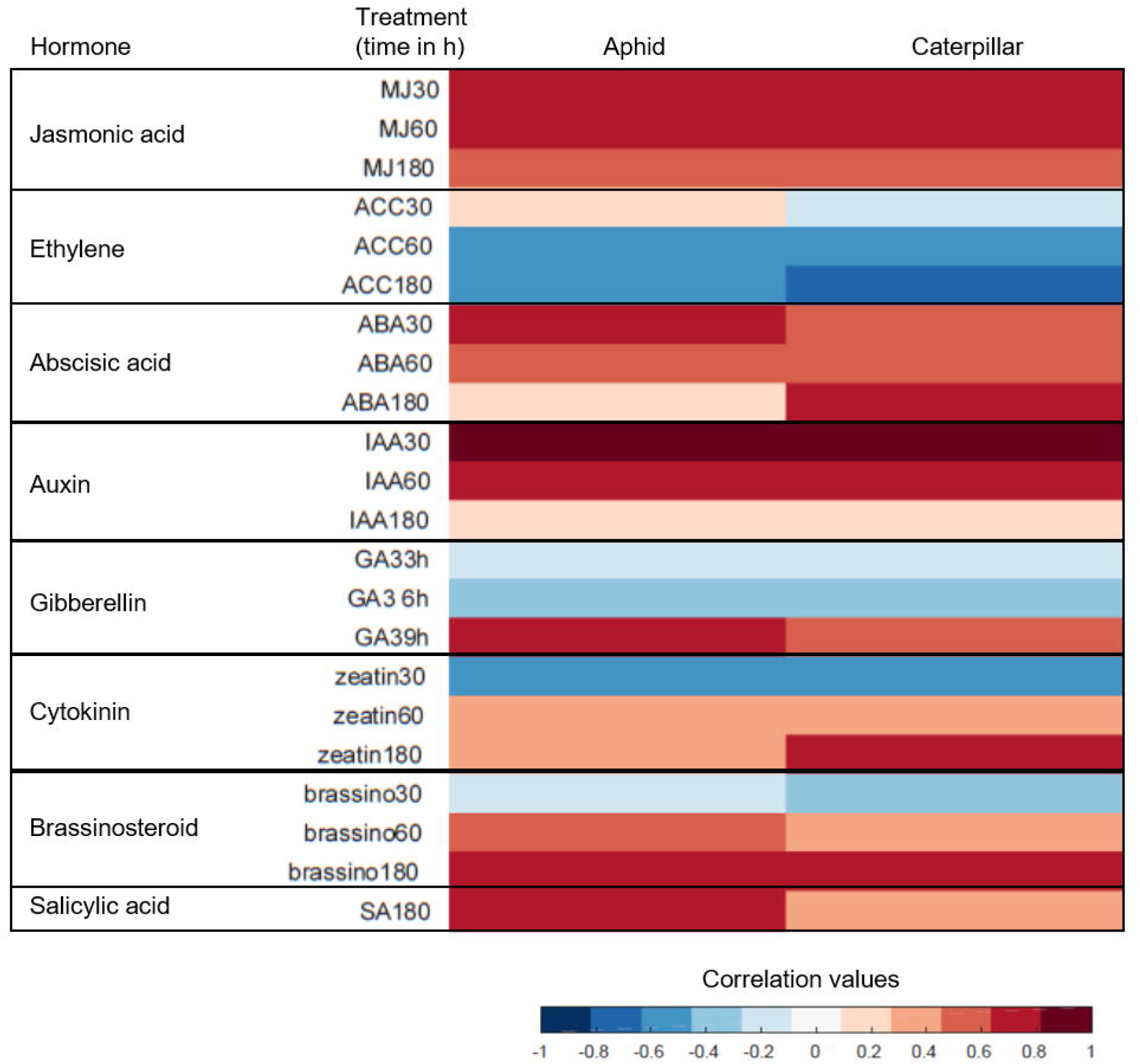
Identification of plant hormone signatures induced by aphids and caterpillar feeding based on transcriptomic data. The analysis was conducted using the Hormonometer program (Oa *et al.*, 2009). Red shading indicates a positive correlation and blue shading indicates a negative correlation between the *S. viridis* herbivore feeding and particular hormone response. The hormones are: MJ, Methyl jasmonate; ACC, 1-aminocyclopropane-1-caroxylic acid (a metabolic precursor of ethylene); ABA, abscisic acid; IAA, indole-3-acetic acid; GA, GA3, gibberelin; BR, brassinosteroid; SA, salicylic acid.

### Serotonin biosynthesis gene identification using rice homologs

To identify the biosynthetic pathway of the secondary metabolites involved in insect defense of *S. viridis*, we focused on indole- and Trp-derived metabolites which have defensive functions in many Gramineae (Grün *et al.*, 2005; Ishihara *et al.*, 2008*b*, 2017; Lu *et al.*, 2018). In particular, we focus on serotonin, which was previously reported as being accumulated in *S. italica* leaves (Kokubo *et al.*, 2016*b*). Serotonin is synthesized in plants via two enzymatic steps: the first is decarboxylation of Trp into tryptamine (Gill *et al.*, 2003; Ishihara *et al.*, 2011; Li *et al.*, 2016; Welford *et al.*, 2016), and the second is oxidation of tryptamine into serotonin ((Fujiwara *et al.*, 2010; Lu *et al.*, 2018); Fig. 1). To characterize the genes, we used both the Plant Metabolic Network database for *Oryza sativa* japonica group (rice) (ORYZACYC 6.0), RiceCyc version 3.3 (http://pathway.gramene.org/ricecyc.html), and comparisons with previous reports (Fujiwara *et al.*, 2010; Dharmawardhana *et al.*, 2013; Lu *et al.*, 2018). For identifying gene homology, we compared the rice and *S. viridis* gene sequences using the Phytozome12 database (https://phytozome.jgi.doe.gov). For the first enzymatic step, we identified ten putative aromatic amino acid decarboxylase (AADC) genes, using all three aromatic amino acids as possible substrates (EC 4.1.1.28) [22]. For the second enzymatic step, we aligned the sequences of the cytochrome P450 monooxygenase CYP71A1, which encodes tryptamine 5-hydroxylase (T5H) in rice, as previously characterized (Fujiwara *et al.*, 2010; Dharmawardhana *et al.*, 2013; Lu *et al.*, 2018). This suggested three putative tryptamine 5-hydroxylase genes (Supplemental Table S8). In Table 2 are presented the putative *TDC* and *T5H* that were significantly modified by either aphid or caterpillar feeding, including five *AADC/TDC* that are induced by both aphid and caterpillar feeding, including *TDC* (Sevir.3G188200; Sevir.9G077100; and Sevir.9G518700), *TDC1* (Sevir.6G066150 and Sevir.6G066200), and T5H (Sevir.5G366500), while the other TDC homologs (Sevir.9G259000 and TDC3 Sevir.8G219500, and the T5Hs Sevir.2G113200 and Sevir.8G219600) were induced only by caterpillars. Interestingly, TDC3-Sevir.8G219500 and T5H-Sevir.8G219600 are located closely together on chromosome 8 (Chr_08:35,580,665..35,583,215 and Chr_08:35,586,149..35,588,652 respectively), suggesting gene clustering (Wisecaver *et al.*, 2017). The Plant Metabolic Network database for *S. viridis* (SVIRIDISCYC 2.0; https://plantcyc.org/databases/sviridiscyc/2.0), suggested that this plant synthesizes benzoxazinoids as the main secondary metabolites. However, expression of the predicted benzoxazinoid genes was not altered by insect feeding and benzoxazinoid defensive metabolites were not detected using HUPLC.

**Table 2.** List of candidate genes from the serotonin biosynthesis in *Setaria viridis*. Genes were annotated using both *S. viridis* and rice data generated from RiceCyc, Plant Metabolic Network database, and Phytozome which revealed the serotonin biosynthetic putative genes: Trp *decarboxylase (TDC)* (EC 4.1.1.28), and tryptamine 5-hydrozylase (T5H), a cytochrome P450. Transcripts that were induced by either aphid or caterpillar feeding are presented. In red, gene expression that were significantly induced. In bold, significant differences relative to untreated control, *P* value ≤ 0.05 (false discovery rate [FDR] adjusted).

| *Setaria viridis*<br>gene ID | Reaction<br>EC | Gene name | Rice gene ID | Log FC<br>aphid/<br>con | *P* value<br>(FDR)<br>aphid/con | Log FC<br>caterpillar<br>/con | *P* value<br>(FDR)<br>caterpillar<br>/con |
| --- | --- | --- | --- | --- | --- | --- | --- |
| Sevir.3G188200 | 4.1.1.28 | Tryptophan<br>decarboxylase<br>(TDC) | LOC_Os05g4<br>3510 | 1.17 | 5.2E-01 | 5.40 | 3.4E-07 |
| Sevir.9G077100 |  |  | LOC_OS10G2<br>3900 | 1.47 | 3.4E-03 | 2.38 | 1.8E-07 |
| Sevir.9G259000 |  |  | LOC_OS10G2<br>6110 | 0.36 | 7.8E-01 | 5.73 | 3.5E-13 |
| Sevir.9G518700 |  |  | LOC_OS10G2<br>3900 | 1.42 | 3.0E-02 | 2.72 | 2.4E-06 |
| Sevir.6G066150 |  | Tryptophan<br>decarboxylase<br>1 (TDC1) | LOC_OS08G0<br>4560 | 2.66 | 2.6E-03 | 4.39 | 2.2E-07 |
| Sevir.6G066200 |  |  | LOC_OS08G0<br>4560 | 2.72 | 2.2E-03 | 4.43 | 1.9E-07 |
| Sevir.8G219500 |  | Tryptophan<br>decarboxylase<br>3 (TDC3) | LOC_OS08G0<br>4560 | -0.49 | 7.5E-01 | 4.78 | 9.9E-06 |
| Sevir.2G113200 | 1.14.13.- | Tryptamine 5-hydroxylase | LOC_Os07g1<br>9210 | 0.87 | 1.6E-01 | 1.59 | 1.8E-03 |
| Sevir.5G366500 |  |  | LOC_Os07g1<br>9210 | 2.10 | 3.1E-04 | 2.93 | 9.5E-08 |
| Sevir.8G219600 |  |  | LOC_Os12g1<br>6720 | -0.96 | 2.9E-01 | 2.78 | 1.1E-04 |

To validate whether these genes are induced by aphid feeding over a different time period, we infested *S. viridis* plants and measured the transcriptomic changes using qRT-PCR. We decided to focus on one gene *TDC1* (Sevir.6G066200), which was significantly overexpressed by both aphid and caterpillar feeding (Table 2). Also, we infested the plants with *Rhopalosiphum padi* (from the same genus as *R. maidis*), which was the only available colony by the time we performed these experiments. As shown in Fig. 4, *TDC1* was significantly induced by *R. padi* aphid feeding after 6 and 96 h, showing 3.55 and 6.27-fold change respectively. This pattern of gene expression at early (6 h) and late infestation (96 h) time points had been confirmed before and indicated plant adjustment (Tzin *et al.*, 2015). This finding supports RNA-seq results that *TDC1* (Sevir.6G066200) transcript is induced by aphid infestation and also demonstrated the pattern of expression over time.

**Fig. 4.**
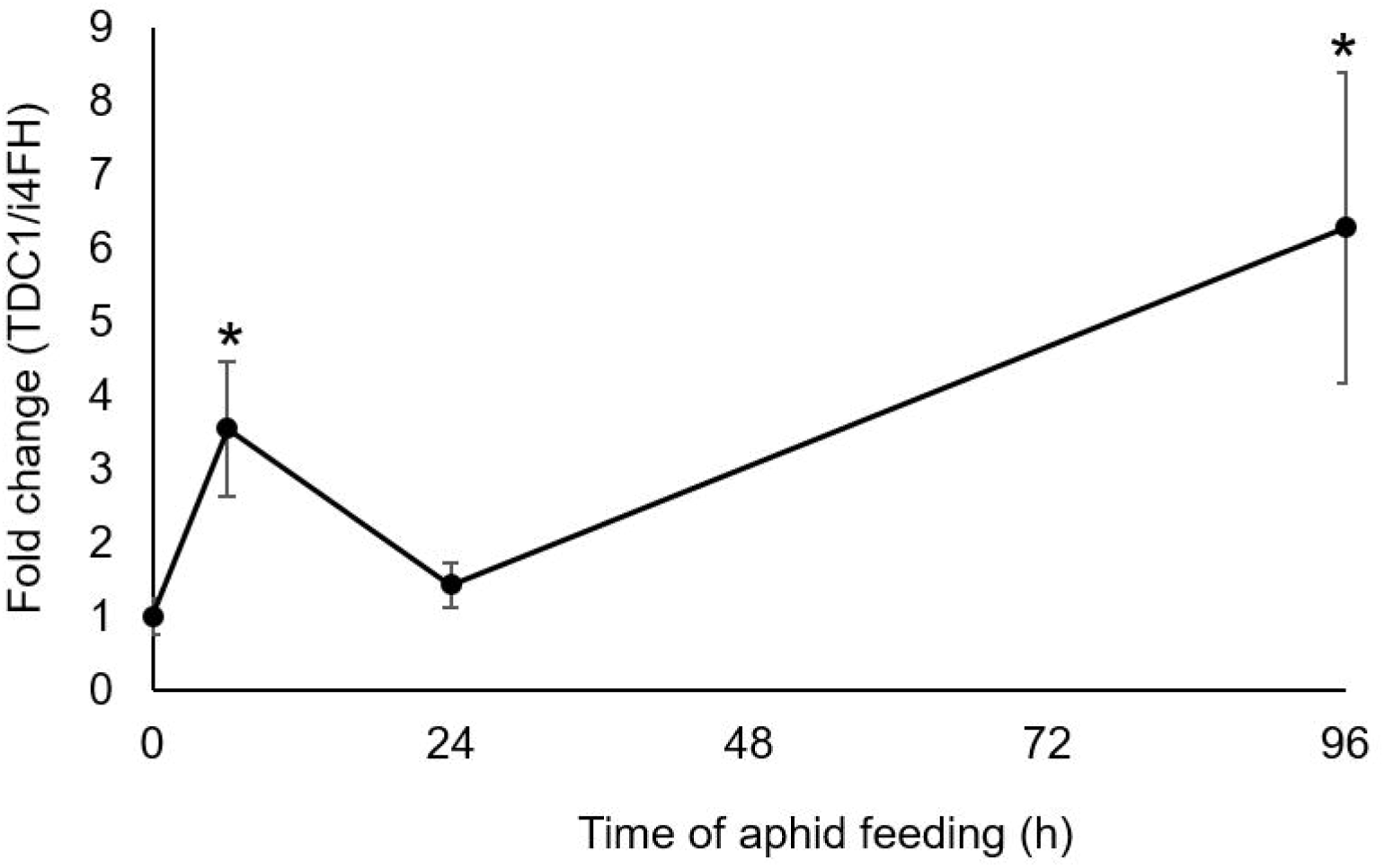
Relative transcriptional levels of the *TDC1* gene (gene ID Sevir.6G066200) in *Setaria viridis* leaves upon *R. padi* aphid infestation. Transcriptional levels were determined at 0, 6, 24 and 96 h after infestation. The asterisk indicates significant differences relative to untreated control (*P* value < 0.05, Student’s *t*-test). Fold change values are of mean transcription (n = 6) with error bars denoting the standard error.

### TDC1 ectopic expression *in planta*

To understand the function of TDC1 *in planta*, its entire coding region was amplified and fused into an expression plasmid using the GoldenBraid system (Sarrion-Perdigones *et al.*, 2014). The gene was fused to an HA-tag, and the *TDC1*-*HA* construct was agroinfiltrated into tobacco leaves. Five days after leaf infiltration, the tobacco leaves were harvested, and expression of the TDC1 protein was quantified by immunoblot. A 63 kDa protein was detected in the extract from transfected tobacco. No band appeared in control with an empty vector, p3α2 (Fig. S1). To validate the function of this protein, metabolites were extracted from tobacco leaves and analyzed by GC-MS analysis. This analysis reveals the appearance of tryptamine in TDC1 transfected leaves, while tryptamine was not detected in the mock-transfected and empty vector leaves (Fig. 5). Overall this suggests that TDC1 can catalyze decarboxylation of Trp to produce tryptamine.

**Fig. 5.**
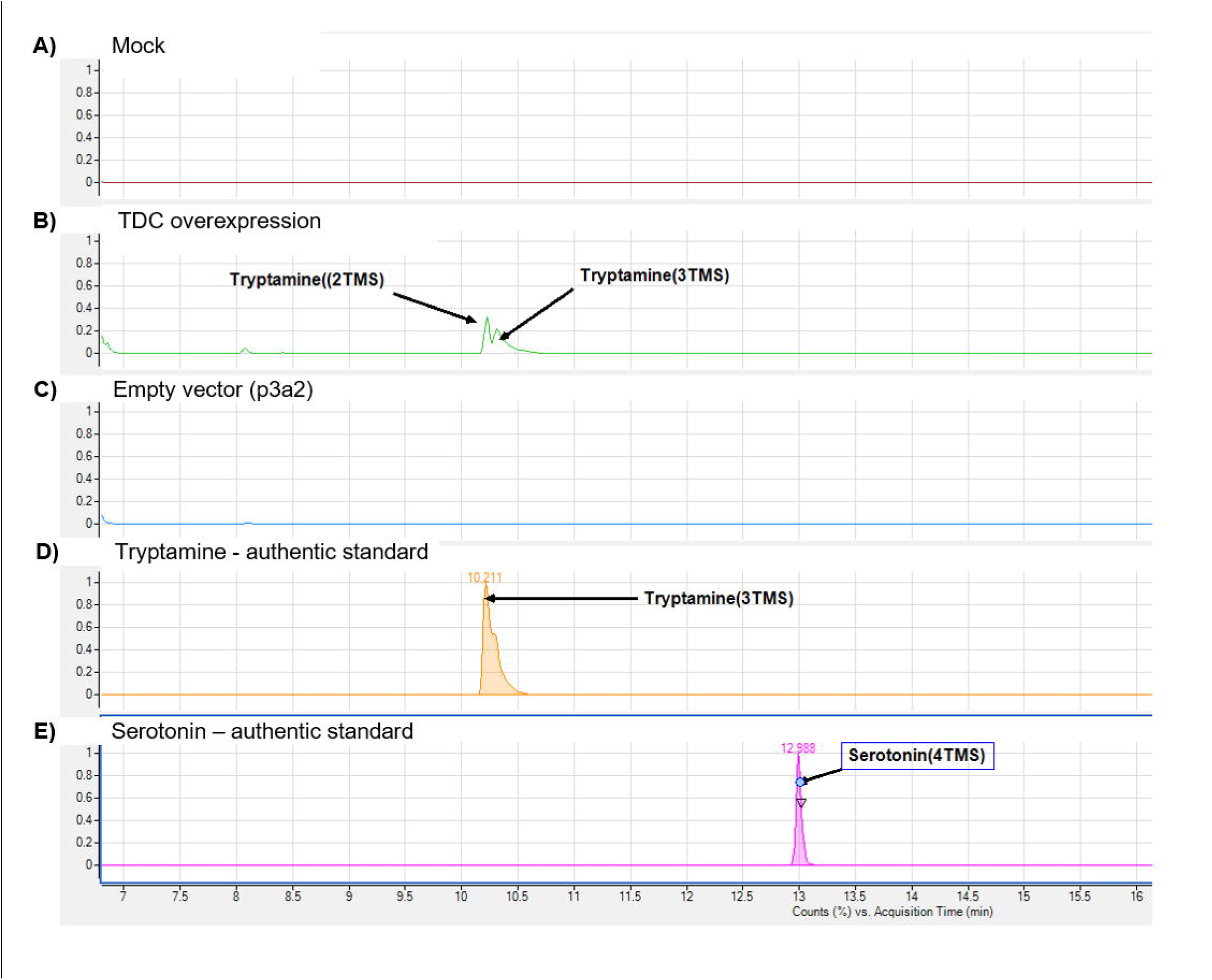
Tryptamine accumulation of TDC1 protein ectopic expression in *Nicotiana benthamiana.* A GC-MS chromatogram of the agroinfiltrated tobacco leaves including A) Mock treatment (infiltrated with buffer only); B) TDC1 ectopic expression and harvesting five days after inoculation; C) Empty vector; D) Tryptamine authentic standard; E) Serotonin authentic standard. The GC-MS chromatograms (A-D) are selected ion monitoring at 174 daltons (molecular mass of tryptamine), and E) is 176 daltons (molecular mass of serotonin).

### Identification of serotonin pathway metabolites in response to aphid feeding

To trigger the biosynthesis of defensive metabolites, we performed a time course experiment of infesting *S. viridis* leaves with *R. padi* aphids for 0, 6, 24, 48 and 96 h. We used GC-MS, to detect the levels of the shikimate-derived metabolites including shikimic acid, Trp, tryptamine and serotonin (Fig. 6). The analysis revealed that shikimic acid is accumulated in the leaves after 24 and 48 h, and serotonin is accumulated after 24 and 96 h. No significant change was detected in the levels of Trp and tryptamine. This is the first indication that serotonin is present in *S. viridis* plants and accumulates in response to aphid feeding. For more information about other primary metabolites including amino acids, nucleosides, organic acids, sugars, and sugar alcohols, that were modified in response to aphid feeding see Supplemental Table S4.

**Fig. 6.**
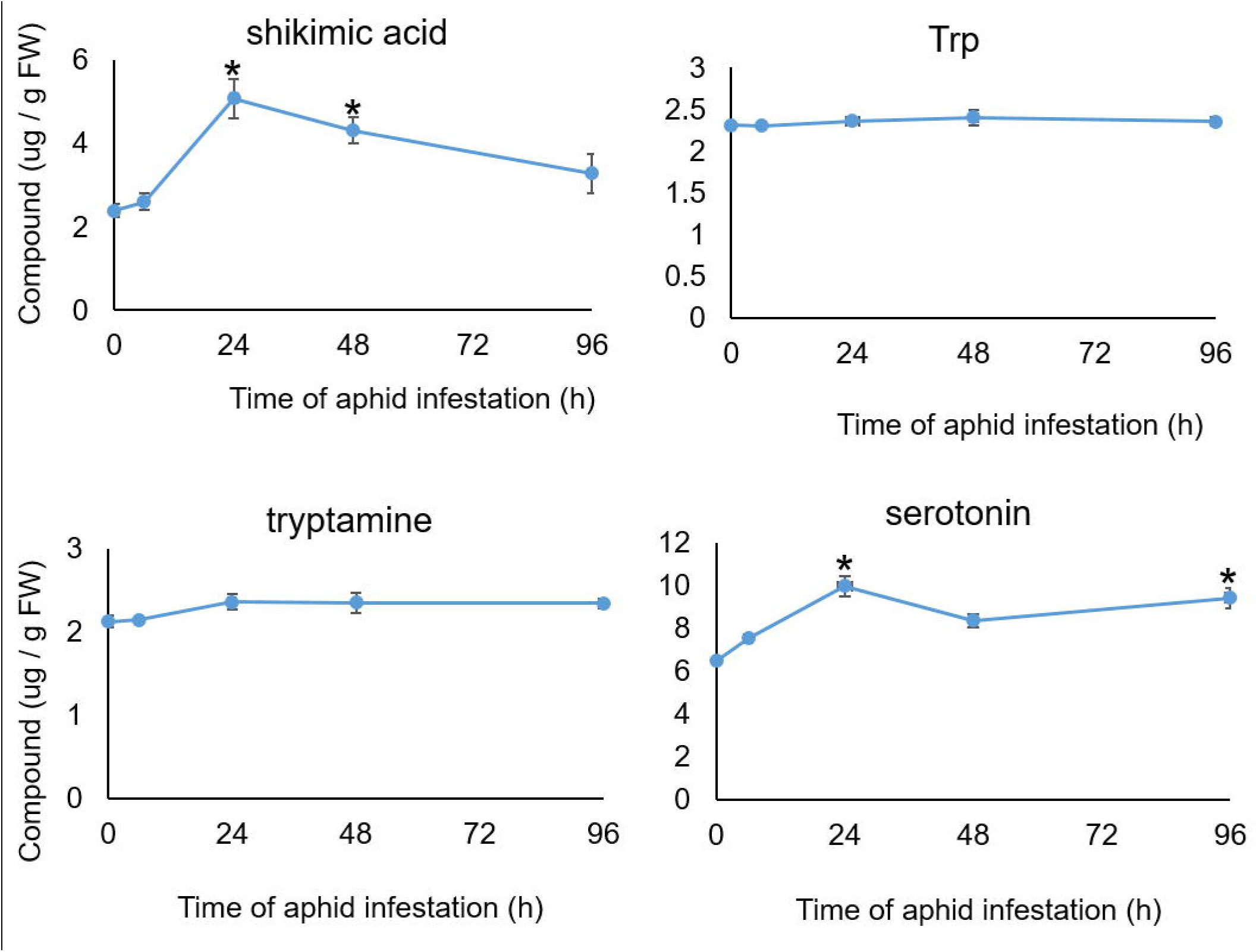
Shikimate-derived metabolite levels after *R. padi* infestation. Metabolite levels were determined at 0, 6, 24 and 96 h after infestation. Asterisks indicate significant differences relative to untreated controls (*P* value < 0.05, Student’s *t*-test). Mean +/− SE of n = 5.

To obtain a more detailed understanding of the contrasting roles of tryptamine and serotonin in aphid resistance, we conducted *in vitro* feeding bioassays with purified compounds. Tryptamine and serotonin were added to an aphid artificial diet at concentrations similar to a previous study (Corcuera, 1984). Trp was used as a control. As shown in Fig. 7, the artificial diet bioassays indicated that, after exposure for two days to the compounds, serotonin was more toxic to *R. padi* than tryptamine and Trp; as 0.28, 0.56, and 1.1 mM serotonin significantly reduced aphid progeny production, and tryptamine and Trp have no effect.

**Fig. 7.**
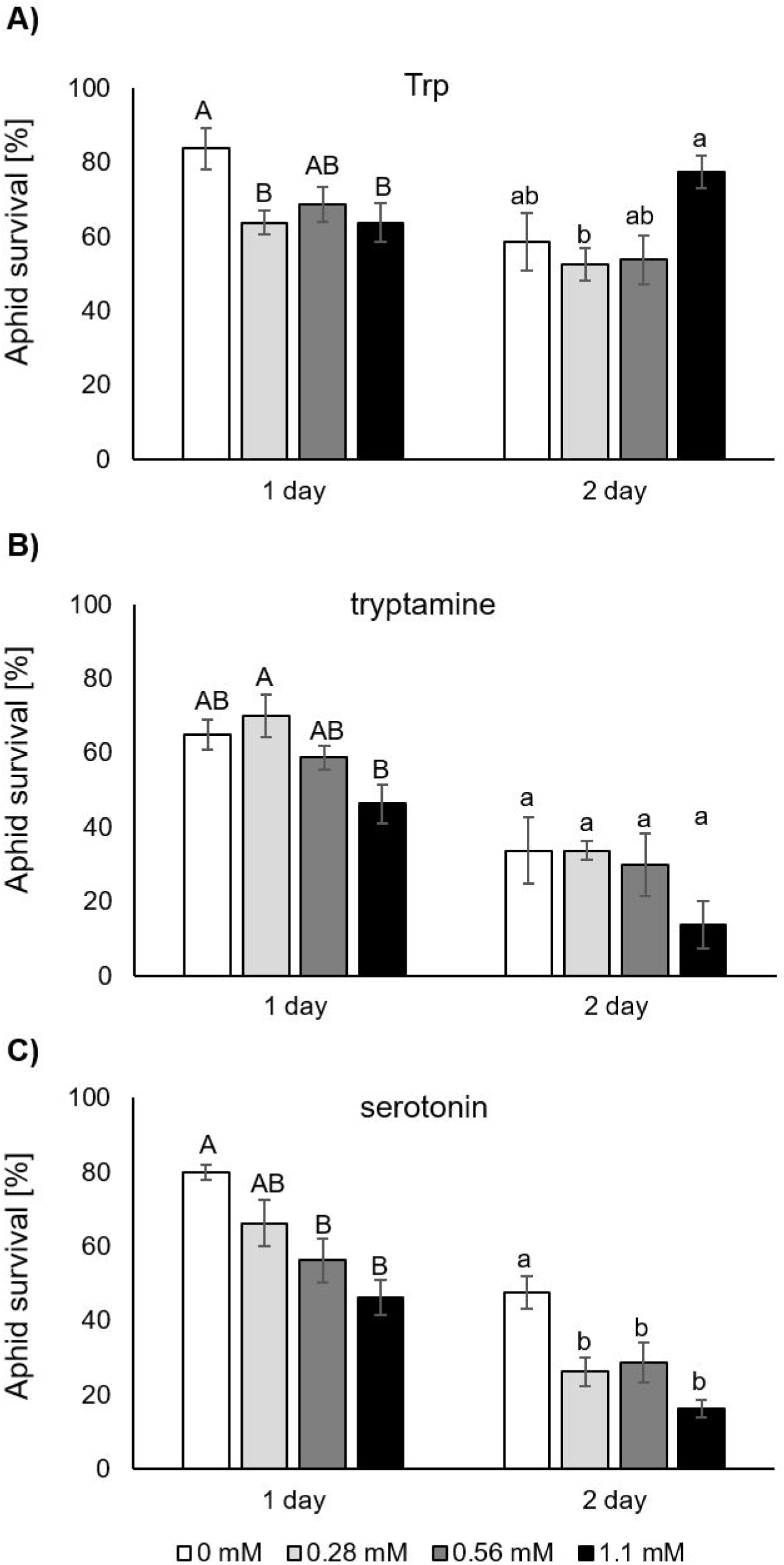
Tryptamine and serotonin reduce aphid performance *in vitro*. Aphids were counted 1 and 2 days following placement on the artificial diet. Aphid survival percentage with a diet supplemented with A) Trp, B) tryptamine or C) serotonin. Bars represent mean values with standard errors (n = 4). Different letters above the bars indicate significant differences (*P* value < 0.05, ANOVA followed by Tukey’s HSD test).

## DISCUSSION

In order to investigate the defense mechanisms of plant responses to herbivore feeding, it is necessary to consider multiple herbivores from different feeding guilds. For example, transcriptomic analysis of *Arabidopsis thaliana* leaves infested by either leaf chewing caterpillars or piercing/sucking aphids identified only a small set of genes that were commonly up- or down-regulated (Appel *et al.*, 2014). However, many comparative studies indicated that chemical defense mechanisms and phytohormones such as jasmonic- and salicylic acid were modified in response to these herbivores (Erb *et al.*, 2012; Kroes *et al.*, 2016, 2017; Pandey *et al.*, 2017). Therefore, we infested *S. viridis* leaves with either *S. exigua* caterpillars or *R. maidis* aphids to unravel the differences in transcriptomic and metabolic effects the two herbivores induce in plants. Comparing the overexpression pathway enrichment and hormonal signature suggested both a unique, as well as a common pattern of gene expression for each herbivore (Fig. 6 and Table 1).

The over-representation pathway enrichment analysis demonstrated that genes associated with amine and polyamine degradation, amino acid metabolism, and biosynthesis of carbohydrates, cell structures, fatty acid, and lipids, hormones, and secondary metabolites were significantly enriched by aphid and caterpillar feeding (Table 1). These observations indicate that there is a shift in primary metabolism and secondary metabolites. The time point experiment of the exposure of *S. viridis* plants to aphid feeding revealed that the metabolic changes occur mainly in the first few hours after infestation (Fig. 6 and Supplemental Table S4).

High throughput methods like RNA-seq analysis are widely used to identify the differential expression of genes (Huang *et al.*, 2018). Hence, RNA-seq was performed to find the differential expression of genes in *S. viridis* leaves following 96 h of herbivore feeding. This analysis showed that putative serotonin biosynthesis genes; *TDC* and *T5H* were upregulated (Table 2). In a previous experiment, it was observed that tryptamine accumulation is preceded by transient induction of TDC, indicating that enhanced Trp production is linked to the formation of serotonin from Trp via tryptamine (Kang *et al.*, 2007; Ishihara *et al.*, 2008*b*). In our study, the expression levels of *TDC1* (Sevir.6G066200) of this gene was upregulated in *S. viridis* infested with either aphids or caterpillars. This suggested that these genes, which are highly expressed in response to insect feeding, are involved in serotonin biosynthesis. The *TDC1* gene was cloned and ectopically expressed in tobacco by the *Agrobacterium*-mediated transformation. The expression and activity of TDC was demonstrated by accumulation of tryptamine. Serotonin was not detected in the *TDC1*-transfected tobacco leaves, which suggests that tobacco lacks this pathway. The function of the candidate cytochrome P450 T5H genes were not shown yet and requires further research.

Trp-related secondary metabolites such as tryptamine and serotonin function as defensive metabolites against pathogens and insect feeding in Panicoideae. For instance, rice plants accumulate these compounds in response to herbivore attacks (Ishihara *et al.*, 2008*b*). The main finding of our research is that *S. viridis* leaves accumulate serotonin 24 and 96 h after aphid infestation (Fig. 6). Serotonin is a neurotransmitter hormone that plays a key role in mood in mammals. However, in plants, it has physiological roles in flowering, morphogenesis and, most importantly, adaptation to environmental changes (Kang *et al.*, 2007). Nevertheless, the effects of serotonin on herbivore growth and feeding behavior is not yet clear. For example, previous experiments using mutants of CYP71A1, which encodes to T5H enzyme, showed that lack of serotonin increased resistance to planthopper (*Nilaparvata lugens*) (Lu *et al.*, 2018), while exposure to the striped stem borer (*Chilo suppressalis*) induced serotonin synthesis in wild type rice plants (Ishihara *et al.*, 2008*a*). This suggests that the function of serotonin in defense is species-specific. Hence, we evaluated the effects of an artificial diet, supplemented with Trp, tryptamine, and serotonin at different concentrations, on aphid survival. The artificial diet experiment showed a reduction in aphid survival at 0.28, 0.56, and 1.1 mM serotonin. The aphid survival rate was reduced on the first day in all three supplemented metabolites. The survival of aphids on both Trp or tryptamine supplemented diet was not significantly modified on the second day. Tryptamine and its derivatives are neuroactive substances that have been found to act as insect oviposition-deterring and antifeedant agent or inhibitor of larval and pupal development (Gill *et al.*, 2003). The artificial diet supplemented with tryptamine also revealed that aphid survival rate was decreased as tryptamine concentration increased, but with no significant difference relative to control at 0.28 and 0.56 mM on the second day. The adverse effects of tryptamine on feeding (Gill and Ellis, 2006) caused the aphids to die from starvation.

## CONCLUSION

Together, the results presented in this research provide new insight into the chemical defense metabolites of *S. viridis*. The transcriptomic analysis elucidated the massive changes that occur after 96 h of insect infestation and provides the first indication that serotonin biosynthesis is involved in the chemical defense mechanisms of this plant species. The induction of serotonin biosynthetic genes, *TDC and T5H*, was also supported by metabolic assays showing that serotonin is induced by aphids. Aphid artificial diet assays demonstrated the potentially toxic nature of this compound. Future research on serotonin biosynthesis induced by aphids and caterpillars will increase the fundamental understanding of the chemical defense mechanisms of crop plants with enhanced resistance to herbivores. In addition, our datasets can be further utilized to discover previously unknown genes and metabolites that function in *S. viridis* and other millets in response to biotic stress.

## SUPPLEMENTARY DATA

**Supplementary Table S1.** RNA-seq raw data and gene annotation.

**Supplementary Table S2.** Primers used for quantitative RT-PCR analysis.

**Supplementary Table S3.** Primers used for the GoldenBraid cloning.

**Supplementary Table S4.** Putative metabolites identified in leaves of *S. viridis* plants infested with *R. padi* aphids analyzed by GC-MS.

**Supplementary Table S5.** LC-MS/MS data from negative and positive ion modes. **Supplementary Table S6.** Gene group generated by the Venn diagrams, compared to rice orthologues.

**Supplementary Table S7.** Orthologous *Arabidopsis* and *S. viridis* genes used for Hormonometer analysis.

**Supplementary Table S8.** Candidate TDC and T5H genes from rice and *S. viridis*. **Supplementary Fig. S1.** TDC1 protein ectopic expression in *Nicotiana benthamina* after five days after agroinfiltration. A) Immunoblots analysis of proteins derived from agroinfiltration *N. benthamina* lines and reacted with anti HA antibodies. An equal amount of proteins was loaded on each lane. B) A loading control showing comparable levels of Ponceau stained proteins in different lanes. M, protein marker.

## ACKNOWLEDGMENTS

We thank Diego Orzaez from the Universidad Politecnica de Valencia, Spain, for providing the GoldenBraid cloning vector collection. We thank Chandrasekhar Kottakota and Moran Oliva for the technical support. Tzin is Sonnenfeldt-Goldman Career Development Chair for Desert Research.

